# Systematic analysis of *Plasmodium* myosins reveals differential expression, localization and function in invasive and proliferative parasite stages

**DOI:** 10.1101/671578

**Authors:** Richard J. Wall, Mohammad Zeeshan, Nicholas J. Katris, Rebecca Limenitakis, Edward Rea, Jessica Stock, Declan Brady, Ross F. Waller, Anthony A. Holder, Rita Tewari

**Author notes:** Corresponding authors: Rita Tewari Richard J. Wall Anthony A. Holder. Richard J. Wall, Anthony A. Holder and Rita Tewari should be considered joint senior authors.

## Abstract

The myosin superfamily comprises of actin-dependent eukaryotic molecular motors important in a variety of cellular functions. Although well studied in many systems, knowledge of their functions in *Plasmodium*, the causative agent of malaria, is restricted. Previously, six myosins were identified in this genus, including three Class XIV myosins found only in Apicomplexa and some Ciliates. The well characterised MyoA, is a class XIV myosin essential for gliding motility and invasion. Here, we characterize all other *Plasmodium* myosins throughout the parasite life cycle and show that they have very diverse patterns of expression and cellular location. MyoB and MyoE, the other two Class XIV myosins, are expressed in all invasive stages, with apical and basal locations, respectively. Gene deletion revealed that MyoE is involved in sporozoite motility, MyoF and MyoK are likely essential in the asexual blood stages, and MyoJ and MyoB are not essential. Both MyoB and its essential light chain (MCL-B) are localised at the apical end of ookinetes but expressed at completely different time points. This work provides a better understanding of the role of actomyosin motors in Apicomplexan parasites, particularly in the motile and invasive stages of *Plasmodium* during sexual and asexual development within the mosquito.

## Introduction

The apicomplexan parasites *Plasmodium* spp., the causative agents of malaria, have three distinct invasive and motile stages (merozoites, ookinetes and sporozoites). Invasion and gliding motility during these distinct stages of the life cycle require the action of an actomyosin motor (Meissner, Schlüter, & Soldati, 2002; Opitz & Soldati, 2002). The myosin superfamily is comprised of molecular motors present during early eukaryotic cell evolution (Foth, Goedecke, & Soldati, 2006; Sebe-Pedros, Grau-Bove, Richards, & Ruiz-Trillo, 2014; Thompson & Langford, 2002). In unicellular parasites they perform a wide variety of cellular functions that require movement (Hartman & Spudich, 2012), including differentiation, host interactions and cell invasion (Meissner et al., 2002). The myosin molecule contains three main domains; the N-terminal head domain, which hydrolyses ATP and binds actin filaments; the neck domain/lever arm, which has an α-helical structure containing up to six IQ motifs; and a tail region, which is required for cargo binding (Thompson & Langford, 2002). The neck domain is known to bind at least one myosin light chain (MLC); proteins with a calmodulin-like domain, which strengthen and extend the lever arm as well as regulate myosin activity (Lowey, Waller, & Trybus, 1993). Although myosins are well studied in many eukaryotes, little is known about the location and function of myosins in the unicellular *Plasmodium* parasite other than their role in motility and host cell invasion.

Within the apicomplexan parasite family, genomic analysis identified six myosin genes in *Plasmodium* and eleven in *Toxoplasma gondii* (*Tg*), the model organism for the study of myosin diversity in apicomplexan parasites (Foth et al., 2006). The phylogenetic analysis, based on sequences of the head domain, allocated each myosin to a specific class within the myosin superfamily, including class XIV, a class that is largely restricted to Apicomplexa. This study also revised some of the myosin nomenclature in the malaria parasite and we have adopted the revised names in this report (Table S1). Only myosin A (MyoA; class XIVa) is well-conserved across Apicomplexa, and a further three myosins, MyoF, MyoJ and MyoK, are common between *Plasmodium* and *Toxoplasma* (Foth et al., 2006). MyoA is by far the most characterised myosin and a vital component of the glideosome motor complex required for gliding motility and host cell invasion (Boucher & Bosch, 2015; Heintzelman, 2015; Meissner et al., 2002). The glideosome is bound to the inner membrane complex (IMC), a system of flattened alveoli located directly beneath the plasma membrane and forming part of the surface pellicle of extracellular parasite stages. MyoA has a crucial, but not essential, role in *Toxoplasma* gliding motility (Egarter et al., 2014; Meissner et al., 2002), but is required for host cell invasion by *Plasmodium* (Sebastian et al., 2012; Siden-Kiamos et al., 2011). MyoA lacks a tail domain but its neck region binds two light chains: myosin tail domain interacting protein (MTIP) and essential light chain (ELC) (Bosch et al., 2006; Green et al., 2017). MTIP links MyoA to the glideosome and the IMC. The class XIVc myosin, MyoB, is the malaria parasite-specific myosin most closely related to *Tg*MyoH, but it lacks a tail domain. It is located at the apical end of the three extracellular invasive stages of *Plasmodium* and has been linked to red blood cell invasion (Chaparro-Olaya et al., 2003; Yusuf et al., 2015). Only one MyoB MLC has been identified, MCL-B, that has an extended N-terminal region and a calmodulin-like domain at its C terminus (Yusuf et al., 2015). The *Toxoplasma*-specific class XVIc myosin, *Tg*MyoH, is located at the apical end (the conoid) of tachyzoites (Graindorge et al., 2016). A conditional gene knockdown approach suggested that this myosin has a role in invasion and is associated with the conoid via its neck and tail domains (Graindorge et al., 2016).

Very little is known about the function or localisation of the four remaining *Plasmodium* myosins. MyoF (class XXII), MyoJ (class VI) and MyoK (class VI) have homologues in *Toxoplasma*. The class XXII *Tg*MyoF localises around the apicoplast during cell division; conditional knockdown results in the loss of the apicoplast in new daughter cells during cell division (Jacot, Daher, & Soldati-Favre, 2013). The class VI *Tg*MyoJ localises as a basal dot in tachyzoites and is implicated in basal complex constriction and rosette formation; gene deletion results in loss of fitness (Frénal et al., 2017). The other class VI myosin, *Tg*MyoK, is located at the centrocone (the spindle pole body) during cell duplication but gene deletion did not reveal a phenotype in the life cycle stages analysed (Frénal et al., 2017). Finally, MyoE (class XIVc) does not have a homologue in *Toxoplasma* and *Tg*MyoE (class XIVb) is phylogenetically distinct from *Plasmodium* MyoE. Other *Toxoplasma* myosins: MyoB, MyoC, MyoD, MyoG, and MyoI have no known essential functions. Given that there are almost twice as many myosins in *Toxoplasma* compared to *Plasmodium* it is likely that different *Toxoplasma* proteins share the same function or have unique functions not required in *Plasmodium*. We therefore systematically characterised all of the myosins in the malaria parasite using the rodent model, *Plasmodium berghei*.

Using GFP-tagging and live cell imaging we showed that all of the class XIV myosins (MyoA, MyoB and MyoE) are expressed at every invasive stage (merozoites, ookinetes and sporozoites). However, the other myosins had very diverse expression and localisation patterns at other unique stages of the life cycle. We found that MyoB is not essential in *Plasmodium*, but its light chain (MLC-B) is. Imaging of ookinete development revealed MLC-B positioned at the apical end early during development, well before MyoB was positioned during the final stages of maturation. Overlap in the localisation of MyoA and the other class XIV myosins suggests a shared function in motility. Finally, functional analysis revealed that MyoE has a role in salivary gland sporozoite motility whereas MyoF and MyoK are likely essential in the asexual blood stage.

## Results

### *Plasmodium* myosins are expressed throughout the life cycle with distinctive patterns of localisation

There are six *P. berghei* myosins, two of these (MyoA and MyoB) have no tail region, and the remainder have a tail, which in the case of MyoF contains five WD40 repeats (Figure 1A). These scaffold-forming repeats construct B-propeller structures allowing the coordination of multi-protein complexes and have an involvement in a wide range of functions. We examined the expression and location of each myosin throughout the life cycle, initially focusing on transcript levels (Figure 1B, Table S2). Quantitative RT-PCR revealed that the class XIV myosins MyoA (PBANKA_0135570), MyoB (PBANKA_0110330) and MyoE (PBANKA_0112200) were strongly transcribed in the invasive stages with an abundance of MyoE transcript in developing merozoites within schizonts (Green et al., 2017; Yusuf et al., 2015). MyoF (PBANKA_1344100) was transcribed at all stages and was second only to MyoA in abundance. In contrast, low levels of transcription of MyoK (PBANKA_0908500) and MyoJ (PBANKA_1444500) could be seen throughout the life cycle (Figure 1B). This distinctive pattern of transcription was largely in agreement with RNAseq data (Otto et al., 2014).

**Figure 1:**
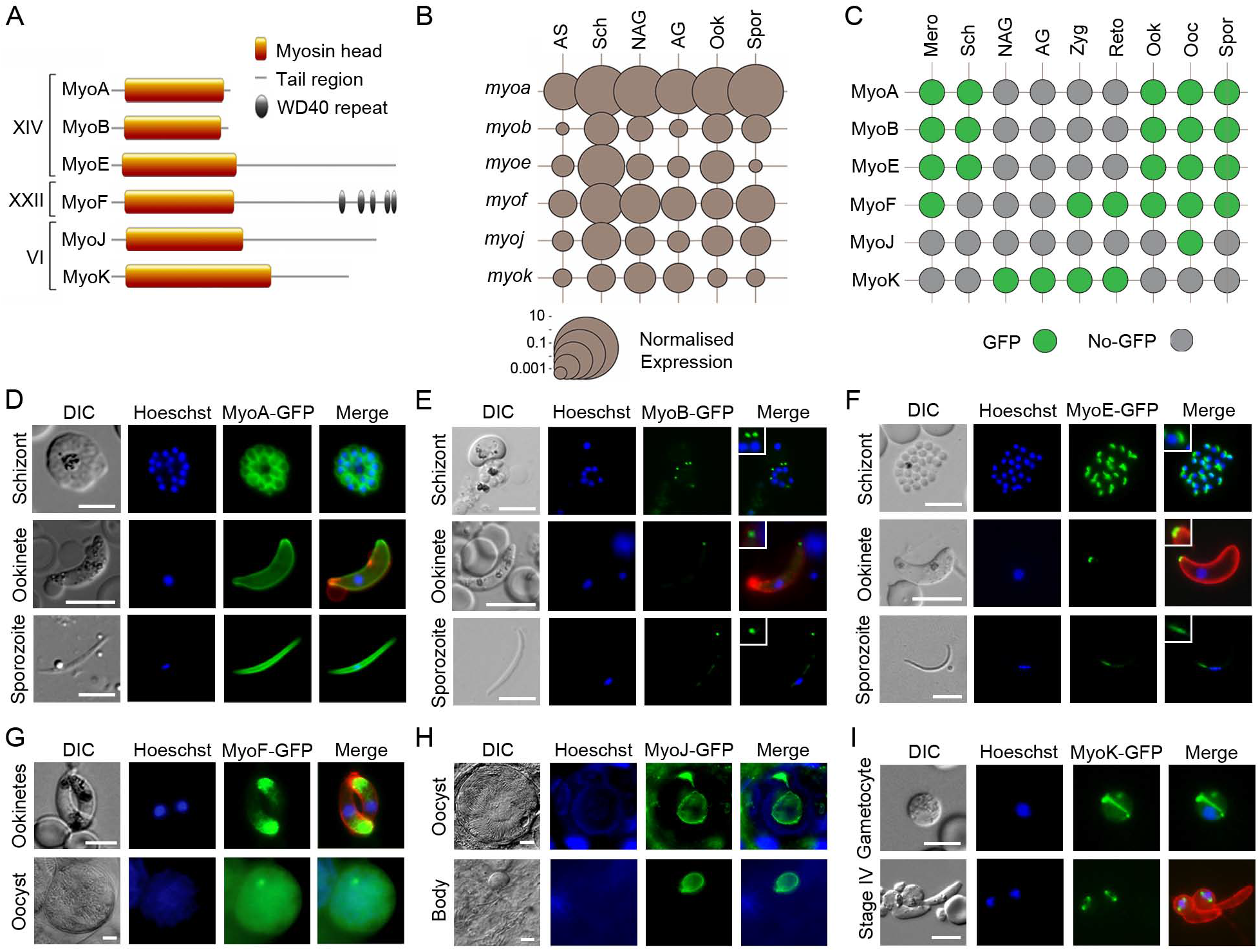
Expression and localisation of *Plasmodium* myosins throughout the life cycle. (A) There are six *Plasmodium* myosins. Two class XIV myosins have a ‘head’ and ‘neck’ region but no tail. The remaining class XIV and class XXII and VI myosins have a ‘tail’, which in the case of MyoF contains WD40 repeat domains. (B) Plot of normalised transcript expression levels for each myosin gene throughout the *Plasmodium* life cycle. RNA was prepared from asexual blood stages (AS), blood stage schizonts (Sch), non-activated gametocytes (NAG), activated gametocytes (AG), ookinetes (Ook) and sporozoites (Spr). Two genes, *arginine-tRNA synthetase* and *hsp70*, were used as controls for normalisation. Each point is the mean of three biological replicates ± SEM. (C) Summary of GFP-tagged myosin expression throughout the life cycle, in merozoites (Mero), schizonts (Sch), non-activated gametocytes (NAG), activated gametocytes (AG), zygotes (Zyg), retort-forms (Reto), ookinetes (Ook), oocysts (Ooc) and sporozoites (Spor). Expression of (D) MyoA-GFP, (E) MyoB-GFP and (F) MyoE-GFP in schizonts, ookinetes and sporozoites using live cell imaging. MyoB and MyoE with higher magnification inset. Expression of (G) MyoF-GFP in ookinetes and oocysts, (H) MyoJ-GFP in an oocyst and residual oocyst body following sporozoite egression and (I) MyoK-GFP in a gametocyte and early (stage IV) retort/ookinete using live cell imaging. Shown are DIC image, Hoechst 33342 (blue); GFP (green); Merge: blue, green and 13.1 (red), a cy3-conjugated antibody recognising P28 (on activated female gametocytes, zygotes and ookinetes only). Size marker = 5 μm.

Next, we investigated the presence and location of the six myosin proteins throughout the life cycle, by adding a C-terminal GFP tag to each via single homologous recombination at the corresponding *myosin* gene locus (Figure S1A). MyoA-GFP and MyoB-GFP have been generated previously (Green et al., 2017; Yusuf et al., 2015). Successful integration was confirmed by diagnostic PCR (Figure S1B). Western blot analysis using an anti-GFP antibody showed MyoF-GFP, MyoJ-GFP and MyoK-GFP were the expected size (Figure S1C). A parasite constitutively expressing GFP and called WT-GFPcon 507 cl1 (WT-GFP) was used as a control. Due to the large size of MyoE-GFP, instead of a western blot, we used immunoprecipitation and mass spectroscopy to confirm that MyoE-GFP was correctly expressed (Figure S1D). No growth phenotype resulted from tagging any of the six myosins (Table S3). We used live cell imaging to detect expression of each protein throughout the life cycle (summarised in Figure 1C) and to examine their location (Figures 1D-1I). The class XIV myosins were detected predominantly in the invasive stages (developing merozoites in schizonts and merozoites, ookinetes, and developing sporozoites within oocysts and sporozoites). As shown previously, MyoA-GFP was associated with the surface pellicle of each invasive stage (Figure 1D) (Green et al., 2017), whereas MyoB-GFP was localised as an apical end dot in merozoites, mature (>20 hr) ookinetes and late stage sporozoites (Figure 1E) (Yusuf et al., 2015). MyoE-GFP was detected as a dot at the basal end of ookinetes and sporozoites, and as a dot in merozoites (Figure 1F). MyoE-GFP was also expressed in liver stages but, as with merozoites, it was not possible to clearly identify the basal end of these stages (Figure S1E). MyoF-GFP was most abundant in the insect stages, particularly ookinetes and oocysts. In mature ookinetes, MyoF-GFP was restricted to the apical end, whereas in oocysts it was more evenly distributed throughout the developing sporozoites (Figure 1G). In contrast, MyoJ-GFP was only observed in mature oocysts, located at the junction between the differentiating sporozoites and the oocyst bodies (Figure 1H). Following oocyst rupture, MyoJ-GFP remained associated with the oocyst body and was not present in sporozoites. Finally, MyoK-GFP was found exclusively associated with the nucleus of gametocytes; appearing as an arc across the cell, and in zygotes/ early ookinetes; as two distinctive dots (Figure 1I). These dots were either absent or only remnants were seen in mature ookinetes.

### Live cell imaging reveals distinct spatio-temporal profiles for each myosin during ookinete development

Ookinete development in the mosquito midgut begins with the growth of a protuberance from the body of the zygote that will ultimately become the apical end of the fully differentiated ookinete. Development of the ookinete transitions through the retort form with further elongation of the developing parasite over a 22 hour period (Janse, Rouwenhorst, Van Der Klooster, Van Der Kaay, & Overdulve, 1985). We showed previously that MyoA-GFP is associated with the pellicle of this protuberance during zygote to ookinete development (Green et al., 2017). MyoA-GFP was only detectable from stage II of ookinete development, initially located throughout the cytoplasm until, after stage V, when it was associated with the pellicle of the growing protuberance and the mature ookinete (Figure 2). MyoB-GFP was not detected until the very last stage of ookinete development, appearing as a dot at the apical end of the parasite (Figure 2). Since *Pf*MLC-B was also expressed at the apical end of merozoites (Yusuf et al., 2015), we next checked its location in zygotes and ookinetes. Generation of a C-terminal GFP tagged line was performed as above (Figure S1A) and validated by integration PCR (Figure S2A). Interestingly, MLC-B-GFP was detected much earlier in ookinete development, visible as an apical dot several hours before MyoB-GFP was observed (Figure S2B). This suggests that MLC-B-GFP is made and positioned, well before its partner, MyoB. MyoE-GFP was detectable as very faint fluorescence in the cytosol until the final stage of differentiation when it was apparent at the basal end of the ookinete (Figure 2). As with MyoB, the localised appearance of MyoE was only seen after the body of the retort was finally reduced to the tapered base of the mature ookinete. MyoF-GFP expression appeared to translocate from the boundary of the stage II ookinetes - where the newly formed ookinete protrudes from the retort - to the apical end of the mature ookinete (Figure 2). As described above, MyoK-GFP was initially detected as a single dot in zygotes, then as two dots in the developing ookinete associated with the nucleus, but the signal almost completely disappeared from mature ookinetes (Figure 2). To resolve the structure of the apical and basal ‘dots’ of MyoB and MyoE (respectively) in mature ookinetes, we used 3D-SIM super resolution microscopy. This showed MyoB-GFP as an apical ring with an aperture in the centre, whereas MyoE-GFP formed a continuous basal ‘cap’ (Figure 3A).

**Figure 2:**
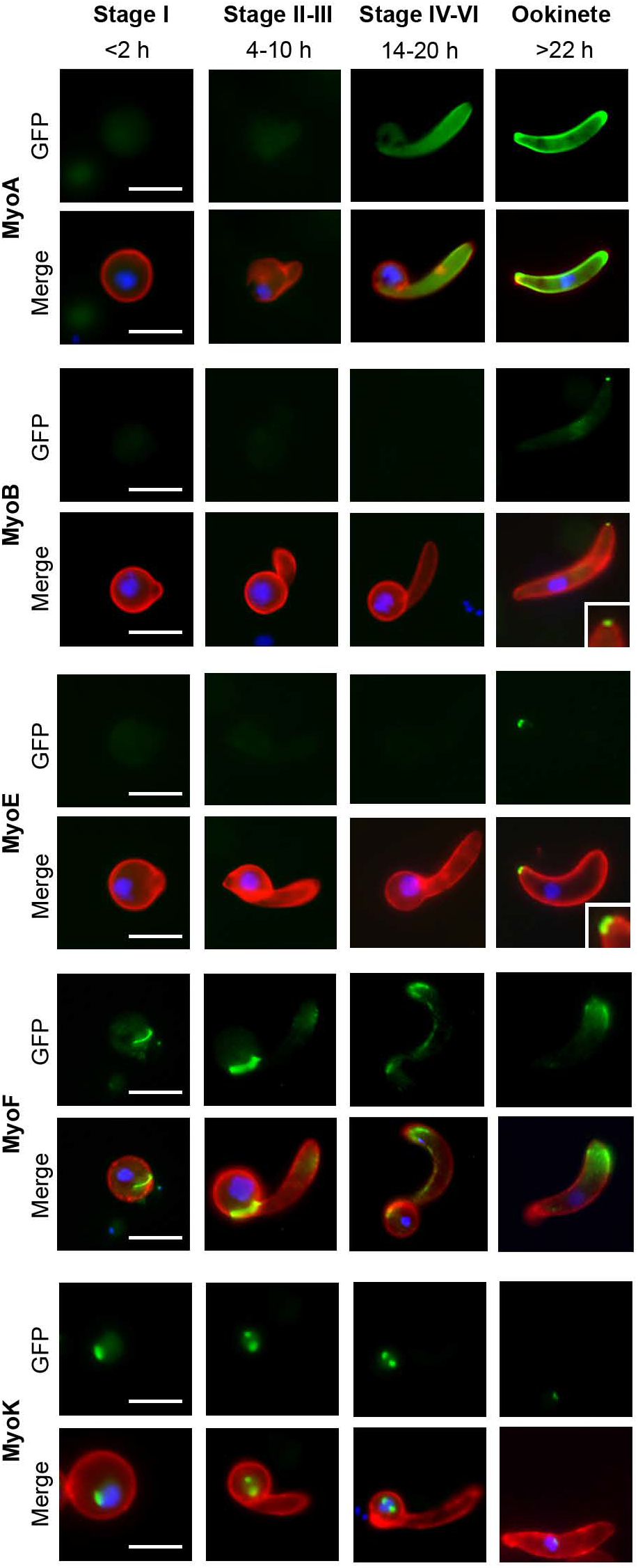
Myosin expression during ookinete development. Localisation and expression of MyoA-GFP, MyoB-GFP, MyoE-GFP, MyoF-GFP and MyoK-GFP during ookinete development (Stage I, <2 h; Stage II-III, 4-10 h; Stages IV-VI, 14-20 h; and mature ookinete, >22 h) using live cell imaging. Top row: GFP (green), bottom row: Merge, Hoechst 33342 (blue); GFP (green); and 13.1(red), a cy3-conjugated antibody recognising P28 on the surface of activated female gametocytes, zygotes and ookinetes. MyoB and MyoE with higher magnification inset. Scale bar = 5 μm.

**Figure 3:**
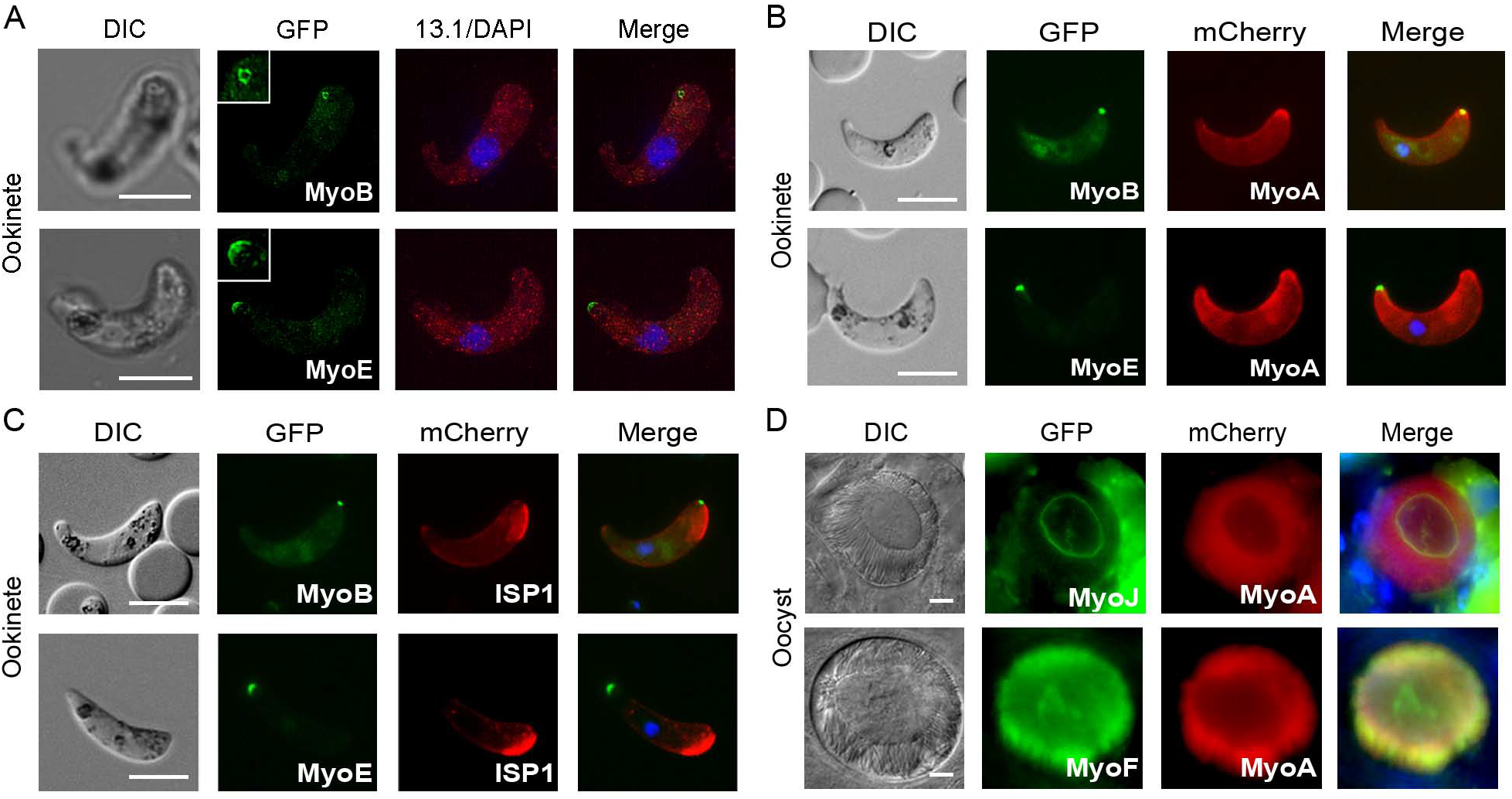
Co-localisation during the life cycle. (A) MyoB-GFP and (B) MyoE-GFP localised by super-resolution microscopy of ookinetes. Shown are DIC image, GFP (green) with higher magnification inset, 13.1(red) and DAPI (blue), and merged image. (C) Co-localisation of MyoB-GFP or MyoE-GFP E with MyoA-mCherry or ISP1-mCherry in ookinetes using live cell imaging. Shown are: DIC image, GFP (green), mCherry (red) and merged image including Hoechst 33342 (blue). (D) Co-localisation of GFP-myosins J and F with mCherry-MyoA in oocysts. Shown are: DIC image, GFP (green), mCherry (red) and merged image including Hoechst 33342 (blue). Scale bar = 5 μm.

### Co-localisation of myosins with MyoA and the apical marker, ISP1

To further study the location of the GFP-tagged myosins, we expressed them in parasite lines that were also expressing either MyoA-mCherry or ISP1-mCherry. ISP1-mCherry is a marker for early ookinete development located and maintained at the protuberance that becomes the apical end of ookinetes (Poulin et al., 2013). The MyoA-mCherry line was generated in this study and validated by integration PCR (Figure S2C). Expression was identical to that seen previously for MyoA-GFP (Figure S2D) (Green et al., 2017). MyoA-mCherry and MyoB-GFP co-localise at the apical end of ookinetes whereas MyoA-mCherry and MyoE-GFP co-localise at the basal end (Figure 3B). These locations were confirmed by MyoB-GFP co-localised with ISP1-mCherry at the apical end, and MyoE-GFP located at the opposite end of the ookinete from ISP1-mCherry (Figure 3C). There was no co-localisation between MyoA-mCherry and MyoJ-GFP, expressed in the sporozoite bodies of oocysts and oocyst body, respectively (Figure 3D). However, the two myosins did overlap at the point where the sporozoites detach from the oocyst body. In contrast, MyoA-mCherry partially co-localised with MyoF-GFP in the sporozoite bodies of oocysts (Figure 3D). Finally, and as seen with ookinetes, MyoA-mCherry also partially co-localises with MyoE-GFP in liver stage schizonts (Figure S2E).

### Systematic gene deletion studies reveal that MyoF, MyoK and MLC-B are essential for asexual blood stages and MyoE has a role in sporozoite motility

To examine further the role of individual myosins, we attempted to delete each of the genes, using double homologous recombination to insert a drug-selectable pyrimethamine resistance cassette in place of the gene (Figure S3A). Previous work had already shown that the MyoA gene cannot be deleted in asexual blood stage parasites (Sebastian et al., 2012; Siden-Kiamos et al., 2011) so we focused on the remaining five myosin genes. MyoB, MyoE and MyoJ genes were successfully deleted, indicating that these proteins are not essential for asexual development in bloodstream parasites. However, despite multiple attempts, we were unable to delete MyoF and MyoK genes, indicating that they, as for MyoA, have an essential role at this stage in the life cycle (See Figure 4A for summary). The three parasite lines, in which individual myosin genes were deleted, were then investigated throughout the life cycle. For each gene, two clones were generated from two independent transfections, and henceforth are known as Δ*MyoB*, Δ*MyoJ* and Δ*MyoE*. Each clone was confirmed to have the correct deletion using diagnostic PCR and Southern blot analyses (Figure S3B-D). In all three lines, male gametogenesis was normal (Figure 4B; Table S4), parasites had a ~70% conversion rate into ookinetes (Figure 4C; Table S4), and all formed oocysts comparable in number to that of WT-GFP parasites (Figure 4D; Table S4). Furthermore, Δ*MyoB* and Δ*MyoJ* also formed a number of both midgut and salivary gland sporozoites comparable with that of the WT-GFP line (Figure 4E; Table S4). In contrast, Δ*MyoE* parasites had a higher number of midgut sporozoites and a reduced number of salivary gland sporozoites compared to the WT-GFP control (Figure 4E). Nevertheless, as with Δ*MyoB* and Δ*MyoJ*, Δ*MyoE* salivary gland sporozoites were able to infect mice as effectively as the control WT-GFP parasites (Figure 4F). However, this reduction in otherwise healthy salivary gland sporozites led us to look at the motility of the Δ*MyoE* sporozoites (Figure 4G-H; Table S4). Motility of sporozoites was analysed using time-lapse microscopy. Despite a significant decrease in salivary gland sporozoites, these sporozoites were found to have normal motility (~55%) compared to WT-GFP (Figure 4G-H, Supplemental Movies S1-6).

**Figure 4:**
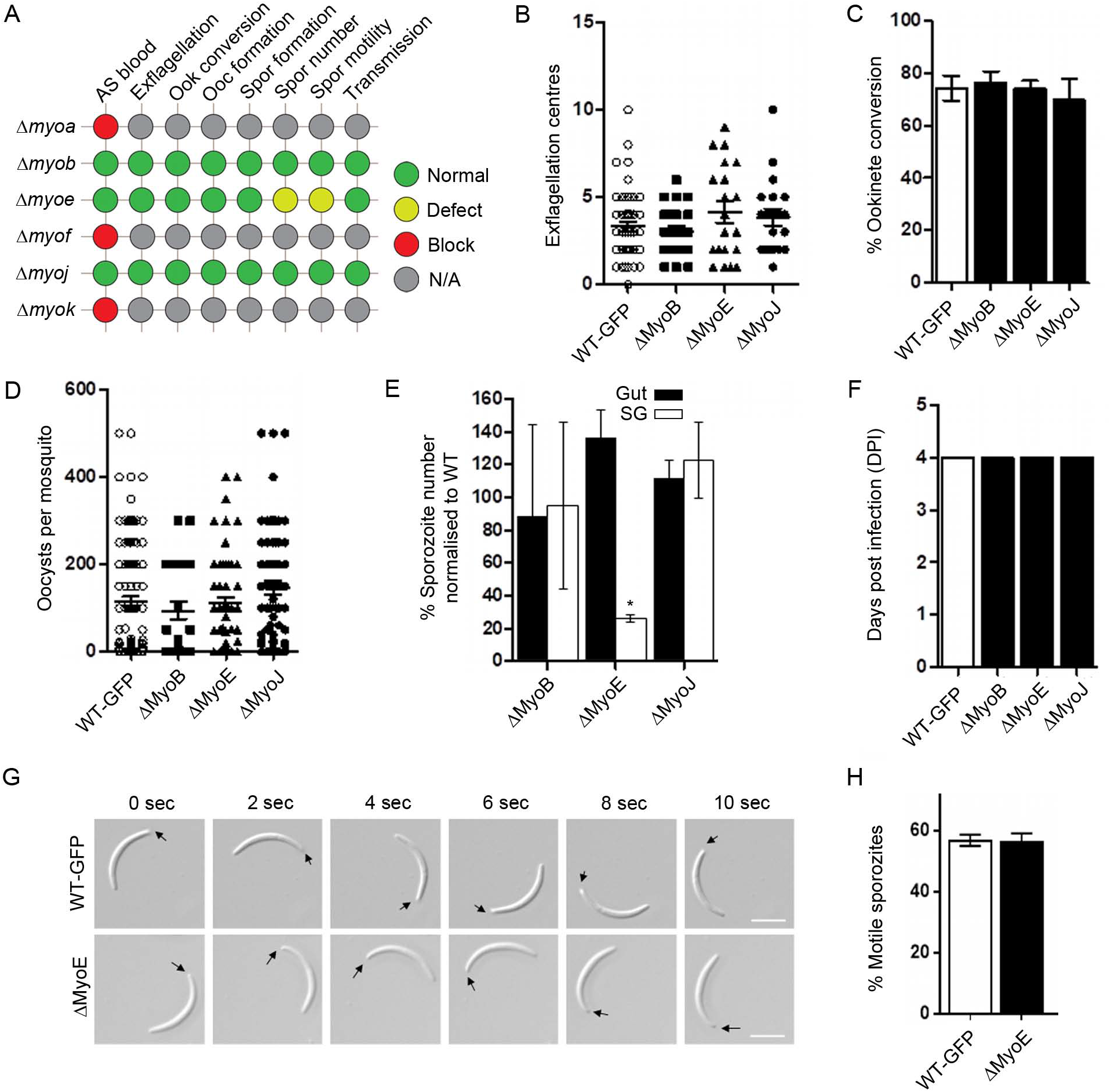
Phenotypic analysis of myosins in *Plasmodium*. (A) Summary of the phenotypic effect of gene deletion in asexual blood stages for all the myosins. For MyoA, MyoF and MyoK, no parasites were recovered indicating that these genes are essential in asexual blood stages, preventing analysis of a phenotype at other stages of the life cycle (red circle; blocked). For MyoB and MyoJ gene deletions, no phenotype was evident at any stage of the life cycle (green circle; normal development). For MyoE gene deletion, a defect was observed in sporozoite number and development (yellow circle). The phenotype was examined in asexual blood stages (AS blood), at exflagellation, ookinete conversion, oocyst formation, sporozoite formation, for sporozoite numbers, for sporozoite motility, and for sporozoite transmission to vertebrate host. (B) Microgametogenesis in gene deletion lines (ΔMyoB, ΔMyoE and ΔMyoJ) in comparison to WT parasite. Measured as the number of exflagellation centres per field, n≥30 from at least three independent experiments with both clones for each line. (C) Percentage ookinete conversion comparing mutant and WT parasites, n≥3 independent experiments consisting of >50 parasites each. (D) Number of oocysts per mosquito midgut (14 days post infection; dpi) for mutant and WT parasites, n≥30. (E) Percentage sporozoite number from midgut (GUT; black bar) and salivary glands (SG; white bar) for mutants normalised to the WT parasite, * *p* ≤ 0.05. n≥3 independent experiments. (F) Bite-back experiments indicating successful transmission of both mutant and WT parasites from mosquito to mouse. Infected mosquitoes were allowed to feed and the mice were monitored for day of patent blood stage parasitaemia (days post infection, dpi) indicative of successful liver and blood stage infection. Example of two independent experiments. (G) Differential Interference contrast (DIC) time-lapse image sequences showing motile WT and ΔMyoE sporozoites isolated from salivary glands. Arrow indicates apical end of sporozoites. Scale bar = 5 μm. (H) Quantitative data for motile sporozoites from salivary glands for WT and ΔMyoE based on two independent experiments.

## Discussion

Myosins have a wide range of motor-related functions and have been well studied in many organisms. However, despite a clear role for acto-myosin-dependant motor functions, almost nothing was known about the myosin repertoire in *Plasmodium*. We present here the first comprehensive functional screen of all myosins in this important pathogen using the rodent malaria parasite model, *Plasmodium berghei*. The different myosins have distinct cellular locations during the life cycle, some of which can be compared directly with those seen in *Toxoplasma*, as exemplified by MyoA. Functional genetic analysis revealed that three myosins are dispensable (MyoB, MyoE and MyoJ) at the asexual blood stage. Previous research has shown that one of the class XIV myosins (MyoA) has an essential role in *Plasmodium* motility and invasion (Sebastian et al., 2012; Siden-Kiamos et al., 2011). Interestingly, we show here that neither of the remaining class XIV myosins (MyoB and MyoE) is essential at any stage in the life cycle. All three class XIV myosins show some level of co-localisation in the three invasive stages suggesting MyoB and MyoE may also have roles in invasion and motility. MyoB has previously been hypothesised to be involved in erythrocyte invasion in *P. falciparum* (Chaparro-Olaya et al., 2003) yet MyoB-disrupted parasites were viable in the asexual blood stages. Although not a direct orthologue of *Pf*MyoB, *Tg*MyoH is also a class XIVc myosin and localises to the apical end as a ring, but in *Toxoplasma* this myosin is essential for gliding motility, host cell invasion and egress (Foth et al., 2006; Graindorge et al., 2016). *Tg*MyoH has three distinct myosin light chains and associates with the microtubules of the conoid, a structure thought to be lacking in *Plasmodium*, although remnants of this structure still survive (Wall et al., 2016). This association occurs through its neck and tail domains and although *Tg*MyoH has a tail domain, this is lacking in *Plasmodium* MyoB. The MyoB light chain, MLC-B, binds to an IQ domain in the MyoB neck region and thereby may act as a tail domain for MyoB (Yusuf et al., 2015). Attempts to disrupt the MLC-B gene were unsuccessful suggesting an essential role during asexual blood stage development. Furthermore, in developing ookinetes, MLC-B is expressed at its apical location up to 12 hours before MyoB suggesting it is the first component of the MyoB-MLC-B complex to be positioned. It is possible that another myosin may substitute for MyoB or, alternatively, MLC-B alone may be sufficient for an unknown essential function in the absence of MyoB. *Tg*MyoH is hypothesised to be involved in conoid function during host cell invasion, namely, contributing to gliding motility where the protruded conoid interacts with the host cell surface before the MyoA glideosome is engaged during invasion (Graindorge et al., 2016). The apparent evolutionary loss or reduction of the conoid from *Plasmodium* suggests that MyoB might not perform exactly the same function. Nevertheless, its light chain, MLC-B, appears to have an essential function in the apical end of the invasive stages. The other class XIV myosin (subclass c; MyoE) was also dispensable for asexual stage development but appeared to have a role in the transfer of sporozoites to the salivary glands. Since motility was not affected, one possibility for the decrease in sporozoites in salivary glands is that MyoE may be involved in sporozoite traversal of the epithelium to the salivary gland duct. Although deletion of the MyoE gene only had a noticeable impact on the transfer of sporozoites to the salivary gland, we cannot exclude the possibility that based on transcript levels, MyoE also has an, albeit dispensable, role at other stages of the life cycle. No MyoE orthologues have been identified in other Apicomplexan parasites suggesting a function unique to *Plasmodium*. However, the class XIVb *Tg*MyoC, although from a different subclass, also has a basal end location and is required for functional motility in *Toxoplasma* (Delbac et al., 2001; Frénal, Marq, Jacot, Polonais, & Soldati-Favre, 2014). *Tg*MyoC is an alternatively spliced version of *Tg*MyoB resulting in a different COOH-terminal tail and distinct localisation. *Tg*MyoC shares several MLCs with *Tg*MyoA and was shown to function within a second glideosome complex. No MLCs have been identified for MyoE in *Plasmodium*, but as with *Toxoplasma*, MyoE may share these MLCs with MyoA or other myosins. Interestingly, *Tg*MyoA, although normally dispensable, is essential to parasite motility in the absence of *Tg*MyoC and, conversely, in the absence of *Tg*MyoA, *Tg*MyoC is essential. Thus, the two myosins (*Tg*MyoA and TgMyoC) can complement each other (Frénal et al., 2014). We observed that *Plasmodium*-specific myosins show distinct polarity: with MyoB at the apical end whereas MyoE was basal in each invasive stage. Furthermore, MyoF also has basal polarity in the early stages of ookinete development and later was found to be at the apical end of the mature stage of ookinete development. Since, by light microscopy, there was clear co-localisation between MyoA and both MyoB and MyoE, we cannot rule out the possibility that MyoA is able to compensate for the loss of either of these proteins in *Plasmodium*, as observed for *Tg*MyoC and *Tg*MyoA. Further studies will be required to investigate this possibility and to fully dissect the function of MyoE during sporozoite transfer to the salivary glands.

Traditionally, class VI myosins work towards the minus end of the F-actin filament (Wells et al., 1999), although whether MyoJ or MyoK follow this rule is not clear from phylogenetic analysis alone (Foth et al., 2006). MyoJ-GFP was observed only in mature oocysts where it localised at the junction between the maturing sporozoites and the oocyst body structure. Previously, residual *Pf*MyoJ was identified in merozoites/schizonts and was suggested to have a putative role in segregating merozoite nuclei (Chaparro-Olaya et al., 2005). Although we also identified MyoJ transcripts at this stage, we did not observe MyoJ-GFP expression. *Tg*MyoJ was shown to be involved in the connection of daughter cells during division; with localisation studies placing *Tg*MyoJ at the basal end of tachyzoites and co-localisation with *Tg*CEN2, suggesting MyoJ might be involved in basal constriction (Frénal et al., 2017). In contrast, MyoK is dispensable in *Toxoplasma* (Frénal et al., 2017) but not in *Plasmodium*. As seen with *Plasmodium* MyoK, *Tg*MyoK appears as two dots representing the centrocones on either side of the nucleus (Frénal et al., 2017). During the sexual stages of *Plasmodium*, micro- and macro-gametes combine to form a 2N zygote which subsequently develops into a 4N ookinete after 22 hr. This complex organisation and replication of genetic material is poorly understood and likely is achieved by the function of a number of components. Centrocones are unique to apicomplexan parasites and are involved in the organisation of the nucleus and the maintenance of genome integrity during cell division (Brooks et al., 2011; Suvorova, Francia, Striepen, & White, 2015). Since we were unable to disrupt MyoF in asexual blood stages, we speculate that this myosin is also performing a similar, essential function at this stage albeit at low protein levels.

An inability to delete MyoF indicates that it has an essential role in asexual blood stage development. MyoF is also essential in *Toxoplasma* tachyzoites, with a role in apicoplast inheritance; in mutants abnormal positioning of this organelle during cell division resulted in failed inheritance and parasite death because the apicoplast performs essential functions (Fichera & Roos, 1997; Jacot et al., 2013). Although *Tg*MyoF is located around the apicoplast in dividing cells, it is present at the apical end of non-dividing parasites (Jacot et al., 2013), as seen for MyoF-GFP in *Plasmodium* ookinetes. Therefore, it is plausible that MyoF may be associated with the apicoplast and have a similar role in *Plasmodium*, however this hypothesis will need to be investigated in future work. MyoF-GFP had an apical location in mature ookinetes, similar but not identical to that of MyoB, although it is abundant throughout ookinete development. MyoF-GFP appeared to translocate from the zygote-ookinete boundary to the apical end of the mature ookinete during differentiation, but whether this is due to movement or *de novo* synthesis is not clear. However, with the potential to form multiprotein complexes via its WD40 repeats, it is possible that MyoF is involved in protein transport during ookinete development. So far, no MLCs or other partner proteins have been identified for the Class VI or XXII myosins in *Apicomplexa*.

Since Apicomplexa have a distinct type of myosin (Class XIV) not found in mammals, these myosins and their MLCs have become the subject of efforts to identify new inhibitors with high selectivity, which may form the basis of new medicines (Douse, Vrielink, Wenlin, Cota, & Tate, 2015; Kumpula & Kursula, 2015; Leung et al., 2014). This possibility highlights the importance of understanding the functions of the other *Plasmodium* myosins. The first comprehensive screen of these myosins described here shows a complex pattern of expression and cellular location, and essential roles for several of the proteins.

## Experimental procedures

### Ethics statement

All animal work has passed an ethical review process and was approved by the United Kingdom Home Office. Work was carried out in accordance with the United Kingdom ‘Animals (Scientific Procedures) Act 1986’ and in compliance with ‘European Directive 86/609/EEC’ for the protection of animals used for experimental purposes under UK Home Office Project Licenses (40/3344 and 30/3248). Sodium pentabarbitol was used for terminal anaesthesia and a combination of ketamine followed by antisedan was used for general anaesthesia. All efforts were made to minimise animal usage and suffering.

### Animals

Six-to-eight week old female Tuck-Ordinary (TO) (Harlan) outbred mice were used for all experiments.

### Protein domain analysis

ScanProsite was used to identify the domains of the myosins (de Castro et al., 2006) and protein domain figures were generated using MyDomains software (Hulo et al., 2008).

### Quantitative RT-PCR

An RNeasy purification kit (Qiagen) was used to isolate total RNA from purified parasites and an RNA-to-cDNA kit (Applied Biosystems) was then used to synthesis cDNA. SYBR green fast master mix (Applied Biosystems) was used for real-time qRT-PCR reactions and data were analysed using an Applied Biosystems 7500fast machine with the following cycling conditions: 95 °C for 20 sec followed by 40 cycles of 95 °C for 3 sec; 60 °C for 30 sec. Gene expression was determined using the Pfaffl method (Pfaffl, 2001) and used *hsp70* and *arginine-tRNA synthetase* as reference genes. Three biological replicates were used for each stage (each with two technical replicates). The primers used are described in Table S5. Statistical analyses (unpaired Student’s t-test) were performed using Excel and Grafit.

### Generation and genotypic analysis of transgenic parasites

For GFP-tagging of each myosin/MLC-B by single homologous recombination, a region of the 3’ end of the gene, but omitting the stop codon, was amplified using the primers listed in Table S5. This was inserted into p277 vector, upstream of the *gfp* sequence, using KpnI and ApaI restriction sites. The p277 vector contains the human *dhfr* cassette, conveying resistance to pyrimethamine. Before transfection, the vector was linearised. The GFP tagging constructs and cell lines for MyoA and MyoB have been reported previously (Green et al., 2017; Yusuf et al., 2015). MyoA-mCherry was constructed using the same primers as described previously (Green et al., 2017). The gene knockout vector for each myosin and MLC-B was constructed using the pBS-DHFR plasmid, which contains polylinker sites flanking a *T. gondii dhfr/ts* expression cassette conveying resistance to pyrimethamine, as described previously (Tewari et al., 2010). PCR primers were used to generate a fragment of 5′ upstream sequence of each gene from genomic DNA, which was inserted into ApaI and HindIII restriction sites upstream of the *dhfr/ts* cassette of pBS-DHFR. A second fragment, generated with PCR primers from the 3′ flanking region of each gene, was then inserted downstream of the *dhfr/ts* cassette using EcoRI and XbaI restriction sites. The linear targeting sequence was released using ApaI/XbaI. *P. berghei* ANKA line 2.34 (for GFP-tagging) or ANKA line 507cl1 (for gene deletion) parasites were transfected by electroporation. Following 2 rounds of pyrimethamine treatment for 4 d each, parasites with successfully integrated constructs were identified using diagnostic PCR. Gene deletion parasites were cloned by limiting dilution to obtain a pure parasite line. For GFP tagged parasites, diagnostic PCR was used to confirm correct integration of the construct and Western blot was used to confirm GFP expression and the correct protein size. For gene deletion parasites, a diagnostic PCR reaction and Southern blot were used to confirm correct integration. These approaches are described previously (Roques et al., 2015).

### Phenotypic analysis

Phenotypic analysis was performed at different stages of the parasite life cycle as previously described (Roques et al., 2015). Briefly, asexual blood stages and gametocytes were analysed using infected blood smears. Gametocyte activation, zygote formation and ookinete conversion rates were analysed using *in vitro* cultures. For mosquito transmission, triplicate sets of 20–60 *Anopheles stephensi* were used. Briefly, exflagellation was examined on day 4 to 5 post-infection. Gametocyte-infected blood was obtained from the tail with a heparinised pipette tip and mixed immediately with 40 μl of ookinete culture medium (RPMI1640 containing 25 mM HEPES, 20% fetal bovine serum, 10 mM sodium bicarbonate, 50 μM xanthurenic acid at pH 7.6). Microgametogenesis was monitored at two different points during mitotic division (8 and 15 minutes post activation [mpa]). Gametocytes were purified and activated in ookinete medium then fixed and processed for immunofluorescence assay (IFA) with antibodies to a range of different markers. Parasites were visualised on a Zeiss AxioImager M2 microscope fitted with an AxioCam ICc1 digital camera (Carl Zeiss, Inc).

### Immunoprecipitation and Mass Spectrometry Analysis

Purified MyoE-GFP schizonts were lysed and incubated with GFP-Trap agarose beads (Chromotek) as described previously (Wall et al., 2018). The sample was then analysed by LC-MS/MS.

### Live cell imaging

Images were captured using a 63x oil immersion objective on a Zeiss AxioImager M2 microscope fitted with an AxioCam ICc1 digital camera and analysed with the AxioVision 4.8.2 software. Ookinetes and zygotes were stained with a Cy3-conjugated mouse monoclonal antibody 13.1 which recognises the P28 protein on the parasite surface (Tewari, Dorin, Moon, Doerig, & Billker, 2005).

### Generation of dual tagged parasite lines

Gametocytes that express the myosin-GFP were mixed with parasites expressing a mCherry-tagged version of either ISP1-mCherry (Poulin et al., 2013) or MyoA-mCherry and incubated at 20 °C for 24 h in ookinete medium. Images were taken as described for live cell imaging.

### Super resolution imaging

To prepare imaging slides, coverslips of thickness No. 1.5H (0.170 ± 0.005 mm) were flamed and smeared with 0.1% polyethylenimine (PEI) solution. Purified ookinetes or salivary gland sporozoites in suspension were fixed with 2% PFA in 1 x PBS and allowed to settle onto PEI-treated coverslips for 20 min and then probed with the indicated antibodies (13.1 mouse monoclonal antibody for ookinetes and CSP antibody for sporozoites) by IFA, stained with DAPI, rinsed in water and mounted in Vectashield on glass slides before sealing with nail varnish. Super-resolution images were acquired using a Deltavision OMX 3D-SIM System V3 BLAZE (Applied Precision) equipped with 3 sCMOS cameras, 405, 488, 592.5 nm diode laser illumination, an Olympus Plan Apo N 60 x 1.42NA oil objective, and standard excitation and emission filter sets. Imaging of each channel was done sequentially using three angles and five phase shifts of the illumination pattern as described previously (Gustafsson et al., 2008). The refractive index of the immersion oil (Cargille) was adjusted to 1.516 to minimize spherical aberrations. Sections were acquired at 0.125 μm z steps. Raw OMX data were reconstructed and channel registered in SoftWoRx software version 6.1.3 (Applied Precision).

### Liver stage imaging

HeLa cells (1 × 10^5^ cells) were seeded in glass-bottomed imaging dishes. Salivary glands of female *A. stephensi* mosquitoes infected with myosin-GFP parasites were isolated and disrupted using a pestle to release sporozoites, which were pipetted gently onto the seeded HeLa cells and incubated at 37 °C in complete minimum Eagle’s medium containing 2.5 μg/ml amphotericin B (PAA), and gassed with 5% CO_2_ in air. Medium was changed 3 hrs after initial infection and once a day thereafter. For live cell imaging, Hoechst 33342 (Molecular Probes) was added to a final concentration of 1 μg/ml, and parasites were imaged at 24, 48, 55 hrs post-infection using a Leica TCS SP8 confocal microscope with the HC PL APO 63x/1.40 oil objective and the Leica Application Suite X software.

### Sporozoite motility assays

Sporozoites were isolated from midgut and salivary glands from mosquitoes infected with WT-GFP and Δ*MyoE* parasites between days 21 and 24 post infection. Isolated sporozoites in RPMI 1640 containing 3% bovine serum albumin (BSA, Fisher Scientific) were pelleted (5 min, 5000 rpm, 4° C). The motility assay was performed as described previously (Moreau et al., 2017). Briefly, a drop (6 μl) of sporozoites was transferred onto a microscope glass slide with a cover slip. Time-lapse videos of sporozoites (1 frame every 1 sec for 100 cycles) were taken using the differential interference contrast settings with a 63X objective lens on a Zeiss AxioImager M2 microscope fitted with an AxioCam ICc1 digital camera and analysed with the AxioVision 4.8.2 software. The assay using matrigel was performed as described previously (Volkmann et al., 2012). A small volume (20 μl) of sporozoites, isolated as above for WT-GFP and Δ*MyoE* parasites were mixed with Matrigel (Corning). The mixture (6 μl) was transferred on a microscope slide with a cover slip and sealed with nail polish. After identifying a field containing sporozoites, time-lapse videos (1 frame every 2 sec for 100 cycles) were taken using the differential interference contrast settings with a 63X objective lens on a Zeiss AxioImager M2 microscope fitted with an AxioCam ICc1 digital camera and analysed with the AxioVision 4.8.2 software.

## Supplemental Figure legends

**Figure S1:**
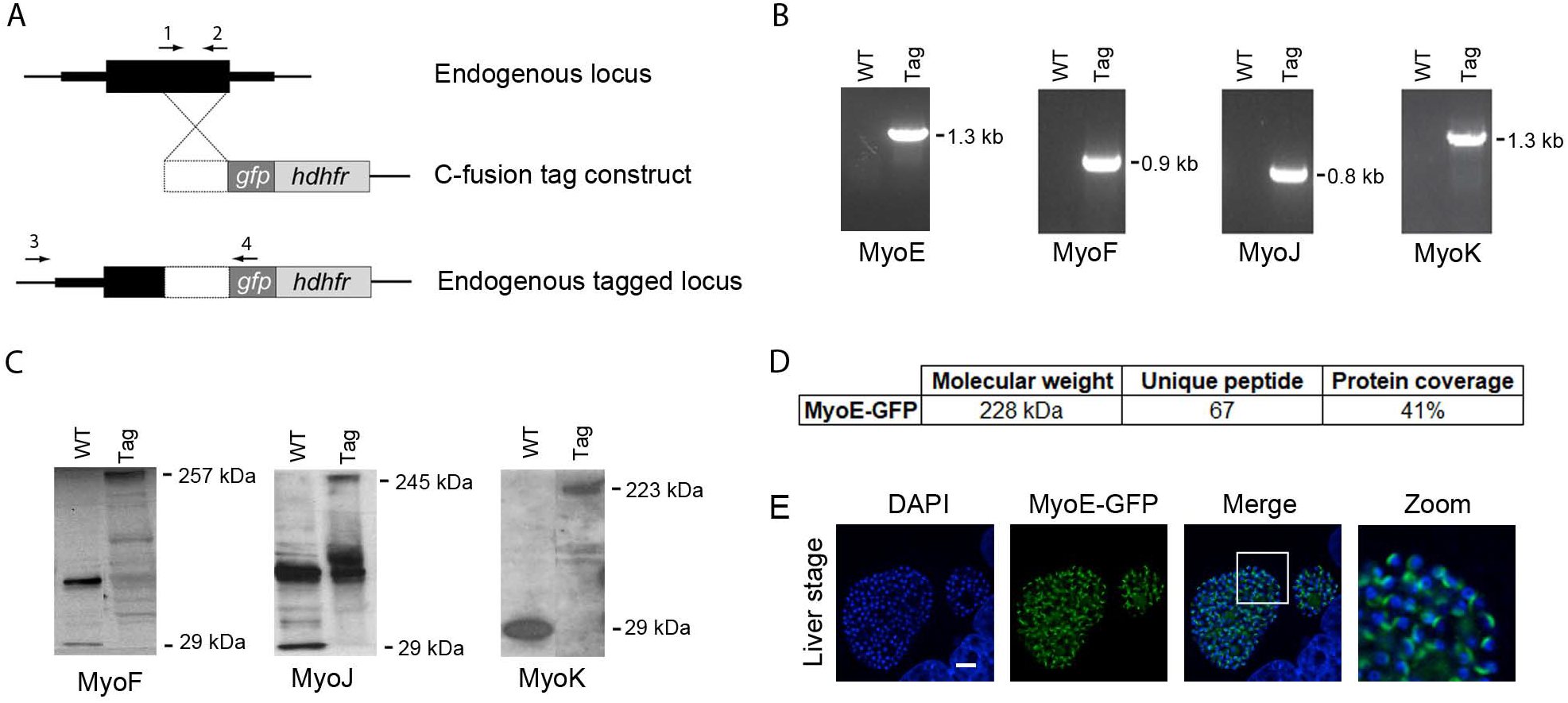
Myosin tag design. (A) Schematic for C-terminal GFP tagging of all the myosins by single crossover integration of the gfp and human dhfr selectable marker at the 3’ end of the gene. The 3’ region of the myosin CDS, immediately upstream of the stop codon, was amplified using primers 1 and 2. (B) PCR analysis to confirm integration for the four myosin tags generated in this project, by amplification of DNA with primers 3 and 4. (C) Western blot analysis of MyoJ-GFP (oocyst) MyoK-GFP (activated gametocytes) and MyoF-GFP (asexual stage) generated in this project, and compared with WT-GFP (29 kDa) using a GFP-specific antibody. (D) Immunoprecipitation of MyoE-GFP with a GFP-specific antibody followed by tryptic digestion and mass spectroscopy identified 67 unique peptides covering 41% of the MyoE protein sequence. (E) Liver stage expression of MyoE-GFP, showing DAPI (blue), GFP (green) and merge (DAPI and GFP) images. The zoom panel is a higher magnification display of the area enclosed by the white box in the merge image. Scale bar = 5 μm.

**Figure S2:**
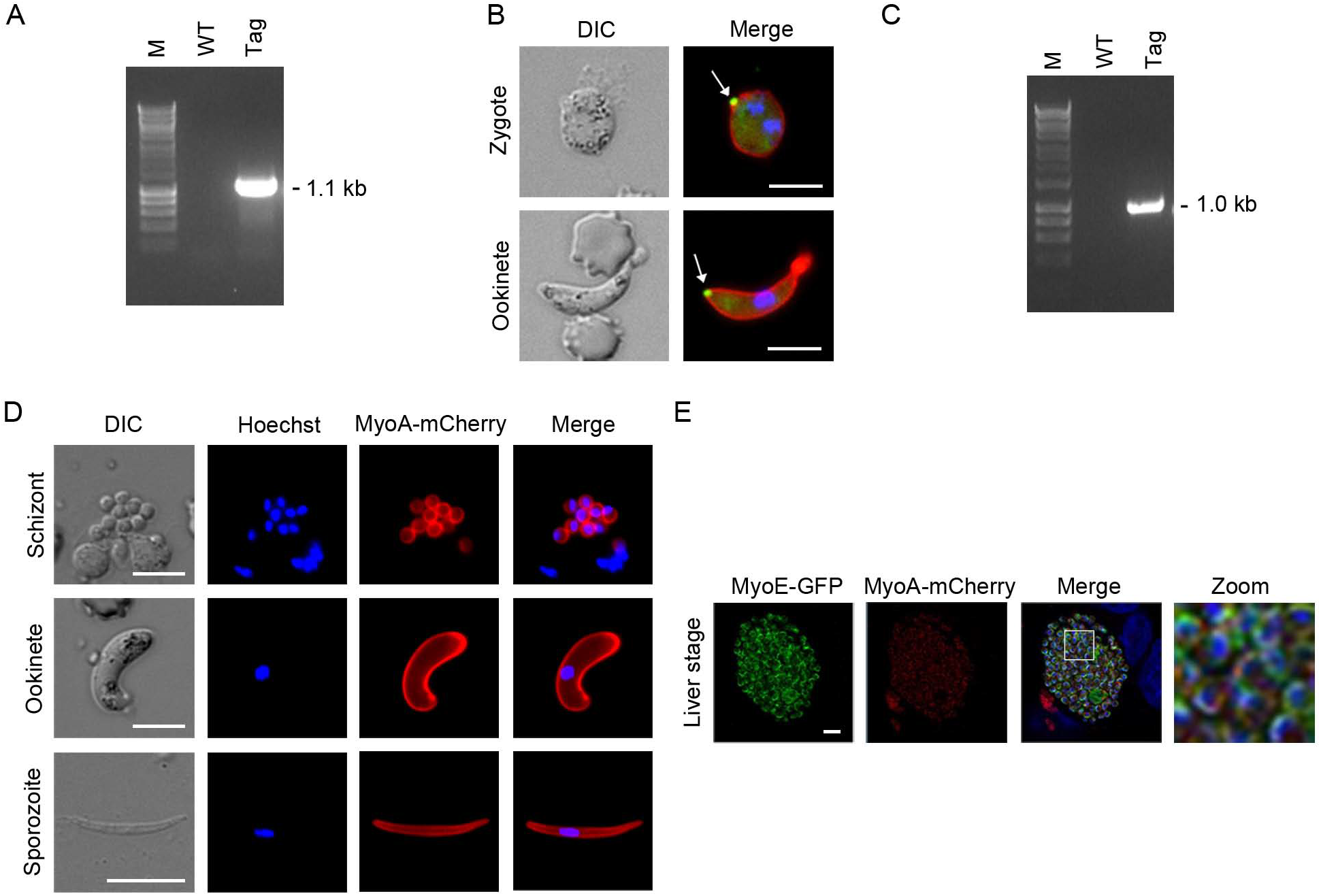
Additional tag validation. (A) Integration PCR analysis of MLC-B-GFP parasite line – based on the same strategy as shown in Figure S1A – using primers 3 and 4. (B) MLC-B expression in both young (<2 hr) retort and ookinete. DIC images are on the left, and on the right are merged: Hoechst 33342 (blue), GFP (green) and 13.1 (red), a cy3-conjugated antibody. White arrow indicates the location of MLC-B protein. (C) Integration PCR validation of MyoA-mCherry parasite line – based on the same strategy as shown in Figure S1A – using primers 3 and 4. (D) Expression of MyoA-mCherry in schizonts, ookinetes and sporozoites (Green et al., 2017). The panels from left to right are DIC, Hoechst 33342 (blue), MyoA-mCherry (red), and merged blue and red. (E) Expression of MyoE-GFP (green), MyoA-mCherry (red) in the liver stage and a merged image including Hoechst stained DNA. A higher magnification image (zoom) is shown. Scale bar = 5 μm.

**Figure S3.**
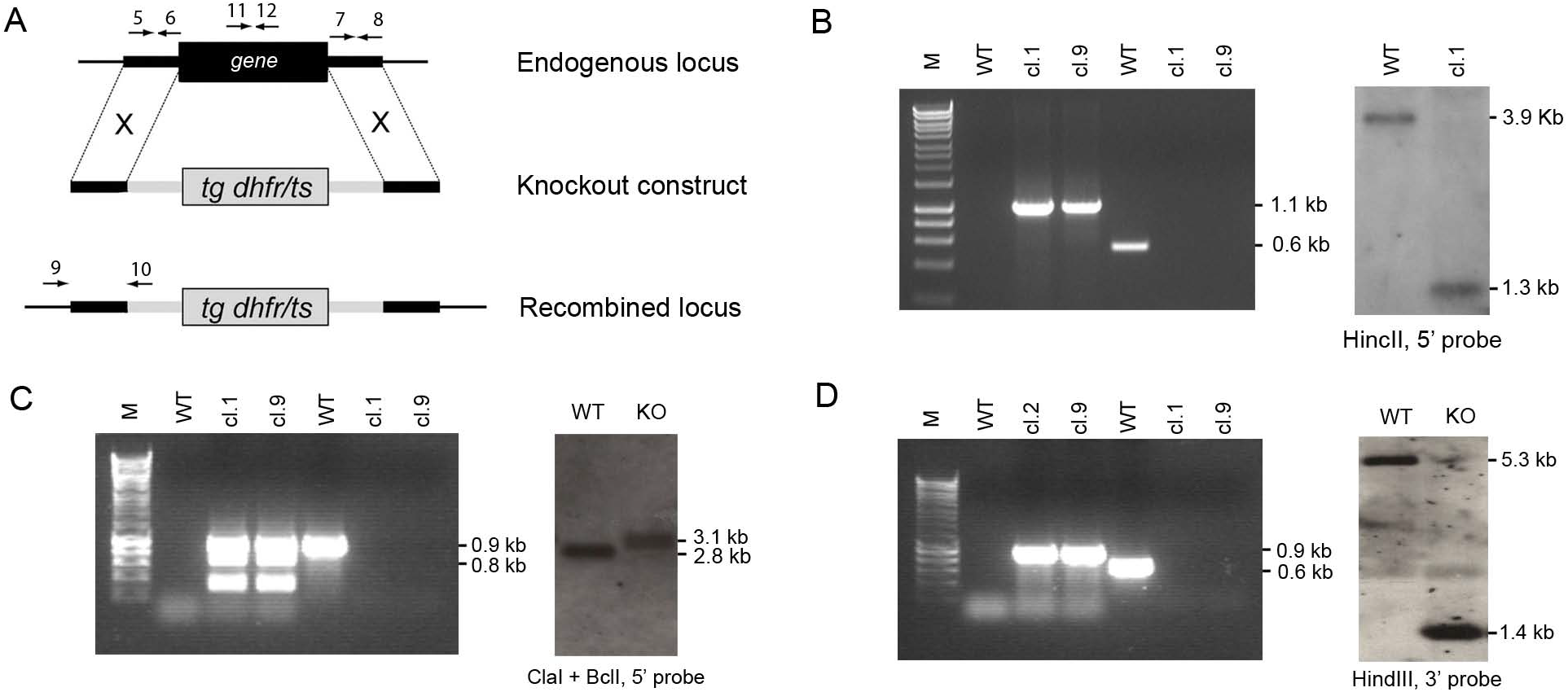
Gene deletion strategy and design. (A) Schematic display of the gene knockout strategy for all the myosins. The drug-selectable DHFR/TS is expressed from the DNA inserted in place of the target gene by double homologous recombination. The 5’ and 3’ region immediately up- and down-stream were amplified using primers 5 and 6 (5’) and primers 7 and 8 (3’). PCR-and Southern blot-analysis diagnostic for successful integration of the selectable marker are shown for (B) MyoB, (C) MyoE and (D) MyoJ. Left side of PCR gels show integration of the construct using primers 9 and 10; Right side of the PCR gels shows successful gene deletion using primers 11 and 12. WT-GFP was used as a control.

## Supplemental Table legends

**Table S1:** Old and new nomenclature and gene accession numbers for *Plasmodium* myosins

**Table S2:** Normalised qPCR data

**Table S3:** Phenotypic analysis of myosin-GFP parasite lines

**Table S4:** Raw data of phenotypic analysis

**Table S5**: Primers used in this study

## Supplemental movies (.AVI files)

**Movie S1**: Gliding motility of WT-GFP salivary gland sporozoite without matrigel

**Movie S2**: Gliding motility of WT-GFP salivary gland sporozoite on matrigel

**Movie S3**: Gliding motility of WT-GFP midgut sporozoite

**Movie S4**: Gliding motility of ΔMyoE salivary gland sporozoite without matrigel

**Movie S5**: Gliding motility of ΔMyoE salivary gland sporozoite on matrigel

**Movie S6**: Gliding motility of ΔMyoE midgut sporozoite

## Acknowledgements

We thank Judith Green for very useful discussions and Julie Rodgers for technical assistance. This project was funded by MRC Investigator Award and MRC project grants to RT [G0900109, G0900278, MR/K011782/1]. RFW was supported by an MRC project grant [MR/M011690/1]. AAH was supported by the Francis Crick Institute which receives its core funding from Cancer Research UK (FC001097), the UK Medical Research Council (FC001097), and the Wellcome Trust (FC001097).

